# Nanoscale chemical tomography reveals organic and inorganic clusters in fossilised dinosaur tooth

**DOI:** 10.64898/2026.09.06.749662

**Authors:** Ya-Lan Hsu, Tim M. Schwarz, Alexander Triccas, Aleksander Kostka, Xinren Chen, Kyuseon Jang, Christina Scheu, Barak Razker, Thomas Tütken, Baptiste Gault, Maïtena Dumont

**Affiliations:** Max Planck Institute for Sustainable Materials, 40237 Düsseldorf, Germany; Center for Interface Dominated High Performance Materials (ZGH), Ruhr University Bochum, 44801 Bochum, Germany; Johannes Gutenberg University, Institute of Geosciences, Johann-Joachim-Becher-Weg 21, Mainz, Germany; Groupe Physique des Matériaux, Université de Rouen, Saint Etienne du Rouvray, Normandie 76800, France

**Keywords:** Atom probe tomography, Biomineralization, Dinosaur teeth, Enamel structure

## Abstract

Mineralized dental tissues are formed through biomineralization processes involving the growth and self-organisation of hydroxylapatite (HAP) nanoscale grains. There remain open questions regarding the way elements such as Mg or Na are incorporated in the organic matter between HAP grains or within the grain structure where they influence the nucleation and growth of HAP. Here, mapping the enamel structure and composition of a well-preserved ca. 150-million-year-old *Giraffatitan brancai* sauropod dinosaur tooth from millimeters down to the near-atomic scale, we reveal the nanoscale accumulation of Mg at HAP grain boundaries, alongside F arising from diagenesis. Within the HAP grains and at HAP grain boundaries organic matter forms clusters that we propose contribute to the fast growth rate of the tooth. Moreover, unexpected Cu_3_As particles are found across the entire enamel structure. These elements are not in the dentine, that exhibits empty tubules, which suggests that they were integrated during the tooth growth itself. Our multiscale analysis provides new information encouraging to reconsider aspects of the biomineralization and fossilization processes.

## Introduction

Fossilized dinosaur teeth offer valuable insights into the diet, physiology, and environmental interactions of these extinct species as tooth enamel is the hardest and consideredly most well preserved hard tissue in a vertebrate body^1,2^. During the early stages of hydroxylapatite (HAP) formation, elements such as Mg, Na, and Fe, as well as C, N, and H, which make up the organic component, play a major role in HAP formation and maturation. Some of these elements are incorporated into the HAP lattice, while others segregate in the intergranular phase between HAP grains^3,4^. In some cases, metallic elements and their distribution in enamel also impact a range of properties, mostly mechanic and acid-resistance – Fe is observed in enamel structure, associated to greater strength and control of the demineralization in shrew, rodents^5,6^ or reptiles^7^ for instance.

In fossils, teeth also retain chemical and structural features that give insights into the biogenic and diagenetic processes^8,9^. Yet, interpretation of the elemental distribution is more challenging, as the surrounding taphonomic setting, including soil/sediment chemistry and environmental conditions as well as diagenetic alteration processes occurring over geological timescales control the fossilization processes^10^. In addition, conventionally-used analysis techniques^11,12^, often fail either to detect elements in low quantity or to resolve their atomic-scale distribution within or in between bioapatite grains^13,14^. The lack of precise, three-dimensional (3D) compositional chemical mapping on the nanometer scale leaves gaps in our fundamental understanding of tooth diagenesis and hampers to distangle postmortem diagenetic alteration from *in-vivo* biomineralization features within fossilized enamel or dentine. Atom probe tomography (APT) could fill this analytical gap as APT provides three-dimensional compositional mapping with nanometer spatial resolution and a chemical sensitivity in the range of tens to hundreds of parts-per-million (ppm) across the periodic table^15^. Since the first report of APT studies to analyze the structure of mineralized tooth structures of mollusk teeth (*Chaetopleura apiculata*)^16^, the technique has increasingly been used to advance the understanding of biomineralization mechanisms and the intimate links between microstructure and properties of enamel structures from different mammals such as humans^17–20^, rodents^3^, and bovids^21^, along with other nonphosphatic biominerals^22–25^, and recently for fossilized teeth^26^.

Here, to explore whether still original *in-vivo* trace element signatures have been retained in the tooth microstructure, we used a correlative multiscale approach involving scanning electron microscopy (SEM), serial sectioning in the dual-beam focused ion beam (SEM-FIB), transmission electron microscopy (TEM), and culminating with APT to analyze a well-preserved fossilized Late Jurassic *Giraffatitan brancai* sauropod dinosaur tooth. APT result reveals the presence Mg-rich and organic-containing nanoscale clusters imaged within and at the boundaries between HAP grains, and the density of these clusters also appears to scale with the size of the HAP grains that grow larger enamel edge to the from the enamel dentine junction (EDJ). In addition, nanoscale copper arsenide (Cu_3_As) particles are seen across from the outer to the inner enamel in between the HAP crystals, and with a morphology that is more rod-like where HAP grains are finer and rounder where the HAP grains were coarser. In light of these new findings, we hypothesised that the clusters contribute to the rapid growth of the teeth by offerering a high density of nucleation sites for the HAP grain growth. We discuss the possible origins of the previously undocumented copper arsenide particles, most likely present within the tooth of the living sauropod through diagenesis, as suggested by the overall preservation of the tooth.

## Results

### Macro- to microstructure analysis of the tooth structure

The fossil tooth of *Giraffatitan brancai*, a sauropod dinosaur from the Late Jurassic, about 150 million years ago, appears well-preserved. This tooth was excavated along with other remains from Tendaguru, Tanzania, during the dinosaur expedition of German researchers led by paleontologist Wilhelm von Branca in 1909–1913^27^. The enamel of the *Giraffatitan brancai* tooth (specimen number MB.R.2181) is around 300 µm thick, three times thinner than the dentine (Figure 1a–b). The dentine shows von Ebner lines, with the most conspicuous ones being 10–25 µm wide (Figure 1c), on par with the report by D’Emic et al.^28^. These daily deposition lines can be used to infer the rates of tooth formation and tooth replacement^29^ which suggests a tooth replacement rate of approximately every 92 days^30–32^.

**Figure 1.**
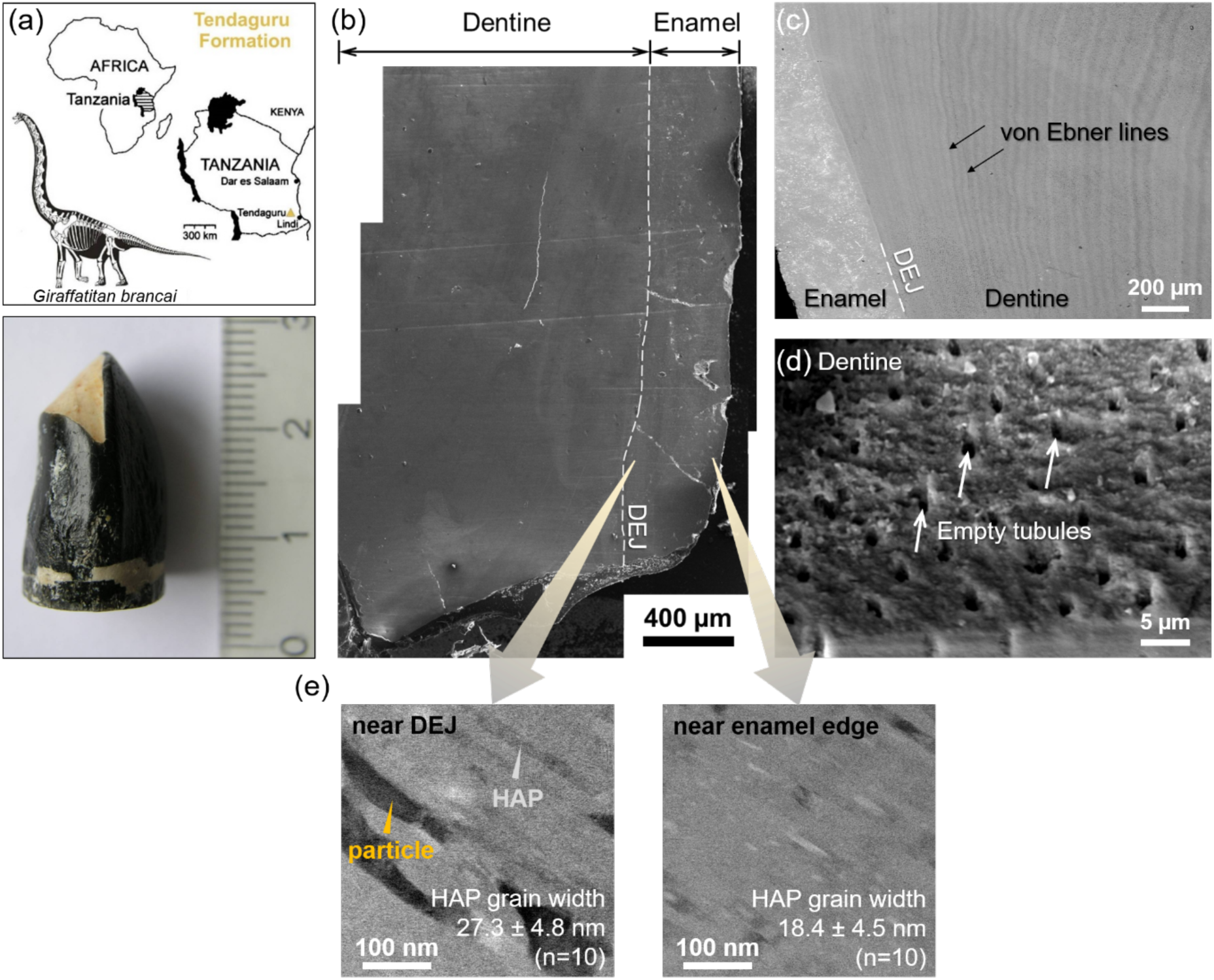
Overview of *Giraffatitan brancai* dinosaur tooth (MB.R.2181) at different spatial scales. (a) Map of the fossil site of Tendaguru, Tanzania, in which the sauropod tooth was excavated from the Late Jurassic (Late Kimmeridgian to Tithonian) sedimentary strata of the Tendaguru Formation. Below the photograph of the tooth is shown (modified from^36^). Note the black colored enamel and the light beige colored dentine. (b) The secondary electron (SE) image of the tooth showing both the dentine and enamel region. (c) A higher magnification image of the enamel and dentine microstructure with clear von Ebner lines in the dentine and enamel dentine junction. (d) The dentine microstructure with empty dentinal tubules indicates the lack of postmortem secondary mineral infillings suggesting a well-preserved fossilized tooth. (e) Bright field TEM images from different regions in enamel (near EDJ and near enamel edge) showing the average width difference of the grains.

The imaged tubules in the dentine structure indicated by white arrows in the SEM image in Figure 1d appear to not be filled with sediment. If, by itself, the absence of secondary mineral infilling within dentinal tubules does not usuallyprovide conclusive evidence of limited diagenetic alteration ^33,34^ it can be considered as one indicator of relatively good tissue preservation. Cowen et al. ^34^, based on comparisons of fossil vertebrate dental remains ranging from the Devonian to the Cretaceous, showed that secondary minerals can infiltrate and precipitate within dentinal tubules and cavities, reflecting fluid-mediated diagenetic alteration. Consequently, the preservation of the original dentinal microstructure and the absence of secondary mineral infilling can be used as morphological indicators supporting limited fluid-mediated diagenesis.This is further supported by TEM images of well-aligned HAP grains in enamel as well as empty preserved dentinal tubules and thin HAP grains structure^35^ in the dentine, as displayed in Figure S3 and Figure S4, respectively.

Bright field (BF) TEM images close to the EDJ in Figure 4e show the assembly of parallel HAP grains aligned throughout the enamel. The specific location where each of the TEM, APT and SEM-FIB tomography analyses were performed is marked (Figure S5). Interestingly, the HAP grains near the EDJ are nearly 10 nm wider on average (over of 10 grains) compared to the HAP grains near the outer enamel edge, which has been reported in humans^37,38^ but never before in dinosaur teeth. Alongside the HAP grains, particles with a darker contrast (Figure 1e, orange arrow) were also observed. These particles are hundreds of nanometers in size, larger than the 2–6 nm gaps that can be found in enamel ^39^, and their detailed characterisation and origins will be discussed in details below.

### APT analysis of the enamel microstructure

We first performed scanning diffraction in the TEM, a technique referred to as 4D-STEM^40^, directly onto an APT specimen prior to its analysis. In 4D-STEM, a diffraction pattern is recorded at each location as the electron beam is scanned over the sample, revealing the distribution of grains within the specimen. This correlative analysis, Figure 2, facilitates the identification of the various features imaged in the subsequent APT measurements. The spatial distribution of the HAP grains is highlighted by a set of red isosurfaces delineating regions containing over 17 at.% of PO_2_^+^ ions, relative to the grain boundary regions. It should be noted that the ionic species detected in APT do not directly reflect the state of the elements within the material^41^.

**Figure 2.**
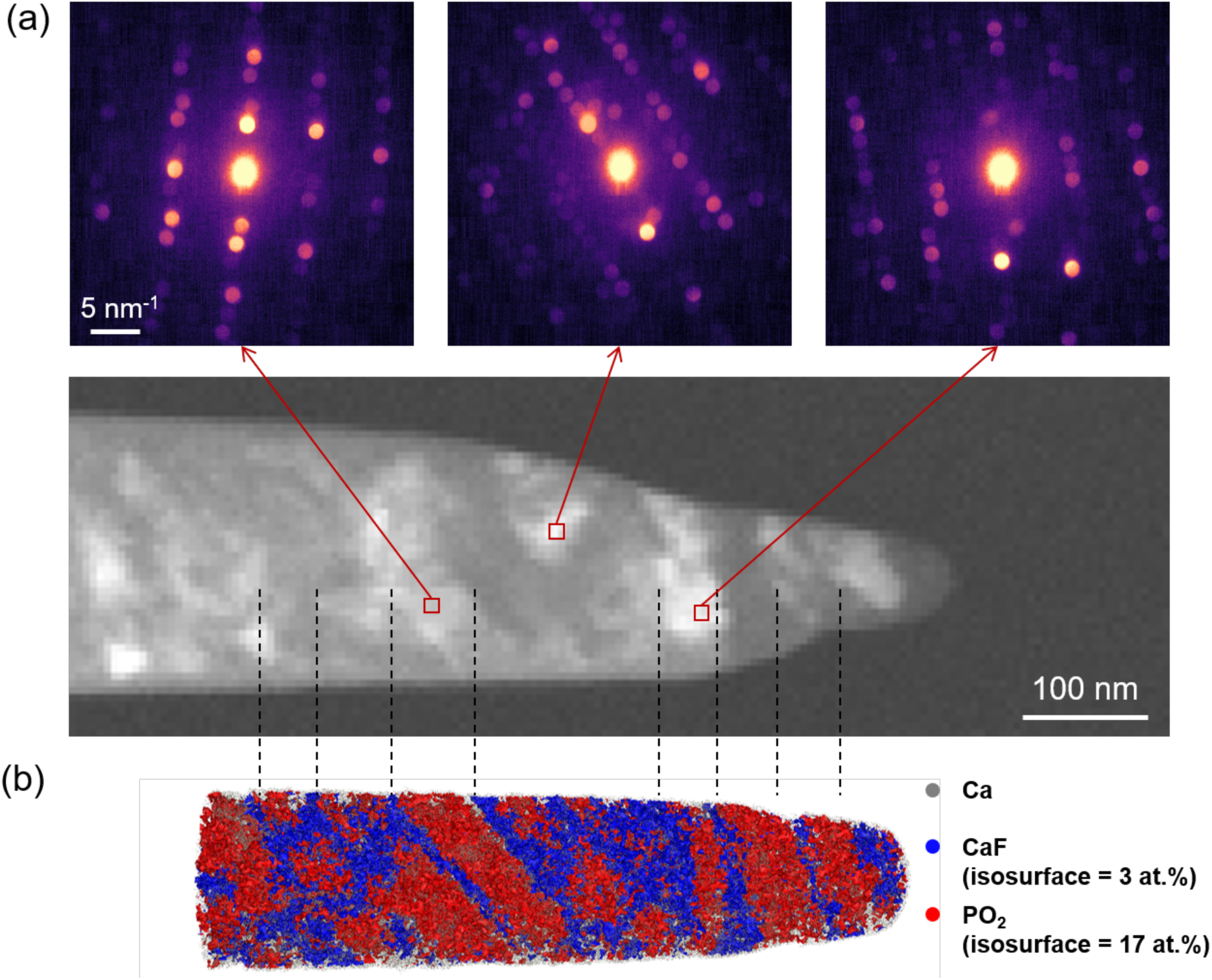
4D-STEM analysis of *Giraffatitan brancai* tooth enamel performed on an APT specimen for direct correlation. (a) 4D-STEM virtual bright field image of the APT sample and the diffraction informations from each red box region. (b) APT reconstruction with an isosurface of 17 at.% PO_2_^+^ and 3 at.% CaF^+^ composition and Ca^2+^ ions dispayed. Note that these ions are related to the APT analysis itself and do not directly reflect the ionic state of these elements within the sample.

This intermediate region between the HAP grains, i.e. the grain boundaries, are evidenced by a set of blue isosurfaces for the CaF^+^ ions. This observation is consistent with DeRocher et al.^19^ who demonstrated that fluoride treatment improved the visualization of the intergranular phases between the HAP grains in human enamel structures and that the CaF^+^ signal was a clear indication of the presence of fluoride in the intergranular phase in APT^19^. Here, however, F most likely originates from fluorine uptake during diagenesis. HAP undergoes chemical changes during the fossilization process, such as the incorporation of trace elements and the exchange of carbonate with the surrounding environment^42,43^, with OH^-^ expected to be replaced by F^-^ ions in the HAP grains, to form the more stable fluorapatite^44,45^. Jang et al. recently reported this in a fossilized dinosaur tooth analysed by APT, where the F measured was across the HAP crystals, and not localized at grain boundaries ^26^. This observation further supports the good conservation state of the analysed sample.

In the analysis of a different specimen displayed in **Error! Reference source not found.**a, isosurfaces of CaF^+^ evidence the alignment of the HAP grains, in agreement with Figure 4e and Figure 2a. At these grain boundaries, F is enriched around three times higher compared to the inner part of the grain (Figure S6). The ionic maps of H_3_O^+^, COH^+^, Mg^2+^ and Na^+^ from another are shown (**Error! Reference source not found.**a), along with close-ups on some of the remarkable features along two orthogonal projections (**Error! Reference source not found.**3b). Top views of region I and region II, as defined in **Error! Reference source not found.**3a, are displayed in **Error! Reference source not found.**3c-d, respectively. Three 1D concentration profiles through the HAP grains calculated along the dashed pink, orange, and green arrows are plotted in **Error! Reference source not found.**3c-1,d-1,d-2, respectively, and key features are indicated by a set of arrows as discussed below.

Pink arrows point to Mg segregation up to 1–1.5 at.% at some grain boundaries (Figure 3c-1), with a Mg content within the HAP grains in the range of 0–0.5 at.%, consistent with reports in human enamel^19^. Mg^2+^ has a smaller ionic radius than Ca^2+^ as well as a greater affinity for water molecules, which limit its incorporation into the enamel grains^46^. It is known that Mg^2+^ ions can replace Ca^2+^ ions during the initial nuclei of amorphous calcium phosphate (ACP), with the lattice mismatch affecting the crystallinity, morphology and lattice parameters, and consequently influencing the HAP grain growth^47,48^. In both regions, H_3_O^+^ and COH^+^ ions, that has been show to originate from organics in biominerals^16,24,49^, agglomerate in 5‒10 nm clusters adjacent to grain boundaries.

**Figure 3:**
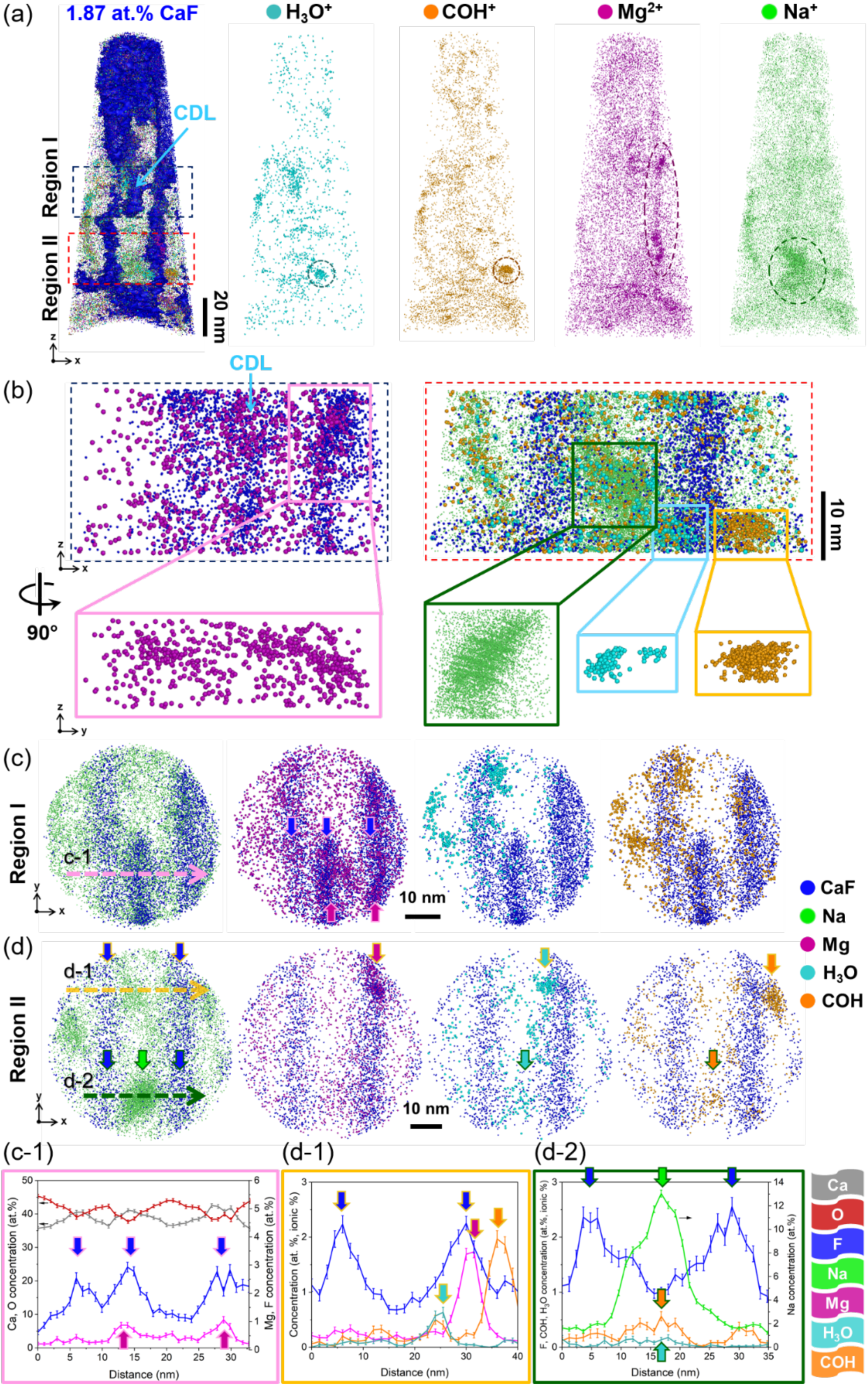
(a) APT 3D reconstruction of the dinosaur tooth enamel structure at the nanometer scale. Isoconcentration surface of 1.87 at.% CaF^+^ was used to visualize the grain boundaries. To observe the distribution of the trace elements and the precipitates, 3D elemental maps of four atoms/ions (H_3_O^+^, COH^+^, Mg^2+^ and Na^+^) are displayed separately. (b) Two regions are extracted from (a) and the four ions are further shown by rotating 90° counterclockwise in z directions. The specimen was prepared from the middle of the enamel (see Figure S5). (c) The cross-section views of APT 3D reconstruction of the enamel structure from region I, and (d) region II. 1D concentration profile measured from (c-1) region I, and (d-1)(d-2) region II.

In region I (**Error! Reference source not found.**c), a F-rich planar feature with the same chemical composition as the grain boundaries was observed in the middle of the HAP grains, together with a segregation of Mg. This could be consistent with the central dark lines (CDL), a planar defect in the middle of the HAP grains, which was often observed in human tooth enamel^18,50–52^, dentine^53^, alveolar bone^54^, and only once in fossil enamel^55^. Many hypotheses have been proposed regarding the structure and chemical composition of the CDL^52,56–59^, with suggested compositional differences based on APT^18^. We can assume that the observed Mg segregation correlates with the CDL, which could however not be observed in the HRTEM images (Figure S3). DeRocher et al.^19^ using X-ray absorption spectroscopy (XAS), electron energy loss spectroscopy (EELS), TEM and APT measurements found an increased Mg content within the core of human enamel HAP grain, which supports our findings.

In enamel (Region II, **Error! Reference source not found.**d) Na is primarily observed within clusters of 2–10 nm wide. There is co-localization with numerous ions usually associated to the fragmentation of organic molecules (e.g., COH^+^, H_3_O^+^)^16,24,49^, within the Na-rich clusters, whereas this is not the case for Mg-rich clusters located at grain boundaries. Organics-containing clusters are often found within the HAP grains also, and similar clusters were observed previously in human enamel^18,60^ and fossilized enamel ^26^ by APT.

Here, these clusters are also Na-rich, and due to solubility reduction during the HAP crystallization process^61^, we can assume that these were originally present during the initial stage of crystallization and might have contributed to the nucleation process. It is proposed that the formation of HAP nuclei is a controlled process, whereby organic matrix proteins (i.e., amelogenin and non-amelogenins such as ameloblastin, enamelins and tuftlein) contribute to the formation of amorphous calcium phosphate (ACP), which gradually transforms into crystalline HAP and becomes organized into highly ordered structures^62–64^. The amelogenin and non-amelogenins are subsequently degraded by proteinases and replaced by minerals^65^. Therefore, near the interface the organic-rich particles found here may be residual degradation products of amelogenin and non-amelogenins as suggested previously^66^, affecting the chemical and mechanical behavior of the intergranular phase e.g., the resistance to crack growth of enamel structure^67^. Indeed, if Mg inhibits crystallization of HAP^47,48,68^, amino-acids can promote it^68^ and carbonates are often implicated in HAP nucleation and increase its solubility, thereby facilitating grain formation^69,70^.

Many similar particles are imaged across all acquired datasests, with further examples displayed in Figure S7 and S8. Therein, we also quantified the content in the primary elements in these clusters, showing a relatively consistent trend in terms of content in organic matter/carbonates, in Na and in Mg. These results are consistent with previous findings in human enamel by APT^18,19^. The incorporation of Na^+^ ions into the HAP lattice requires charge compensation to maintain structural integrity, which is typically achieved by incorporating anions in the sublattice, such as CO_3_^2-^ ions^71^. Figure 3d-2 shows an increase in the COH^+^ signal together with Na^+^, indicating co-segregation of these two ions. In addition, in Figure S9 we report on the volume density of the organic clusters COH in analyses performed near the EDJ and near the enamel edge.

### Copper-arsenic across the enamel

The dark contrast of particles in Figure 1(e) suggests an alternate elemental composition beyond the HAP grains within the tooth. Secondary electron imaging in the SEM in Figure 4a reveals a dense distribution of brightly-imaging particles in the enamel, indicative of a high concentration of heavy elements, and as such consistent with the dark contrast in bright field TEM. No such features are visible and present in the dentine. A TEM lamella containing some of these particles was prepared. In the high-angle annular dark-field (HAADF) micrograph shown in Figure 4b, particles appear with a brighter contrast. Since the contrast scales with the atomic number (Z), these particles contain a high concentration of heavy elements (i.e., Cu and As or Pb) compared to the HAP grains (Ca). The indexing of the selected area electron diffraction (SAED) pattern, from one of these particles, indicates a hexagonal phase (*P*6_3_*cm*, No. 185)^72^ along the [010] zone axis. Energy-dispersive X-ray spectroscopy (EDS) mapping in the SEM (Figure S10) and TEM, Figure 4b, indicate that the particles contain Cu as well as As, Mg or Pb which cannot be discriminated because of overlaps in the EDS spectrum lines (Mg Kα=1.254 keV, As Lα=1.282 keV, and Pb Lα=10.551 keV, As Kα=10.544 keV), see Figure S10.

**Figure 4:**
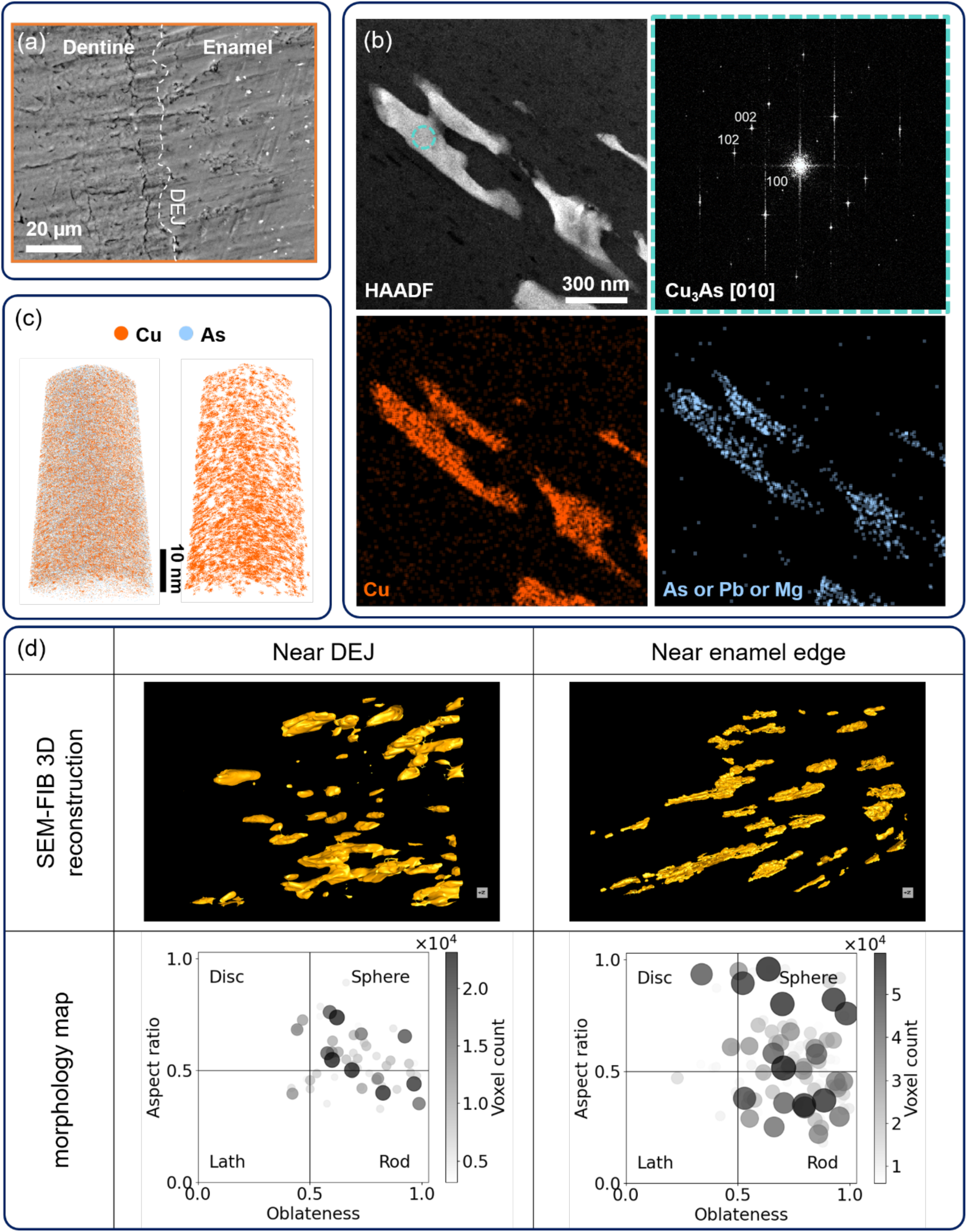
Characterisation of copper arsenide in the enamel across scales. (a) SE images of the dentine and enamel area of the *Giraffatitan brancai* tooth near the EDJ. (b) TEM HAADF image and corresponding TEM-EDS elemental mapping, and the fast fourier transform (FFT) pattern from the teal circle region in the bright contrast in TEM HAADF image. (c) APT 3D reconstruction of the specimen lift-outed from the bright contrast region. (d) SEM-FIB tomography 3D reconstruction and morphology map of the Cu_3_As particles from different enamel regions.

The APT measurement confirms that these are particles consisting of Cu and As, with no Pb or Mg (Figure S11), with a Cu:As ratio near 3:1. However, a precise quantification is difficult due to the field evaporation behavior of As^73^ leading to possible overlaps of different As peaks with the same mass (e.g. As_2_^+^ and As_4_^2+^) in the mass spectrum (Figure S11). The Cu-As phase diagram reports an equilibrium ordered phase, γ (Cu_3_As) copper arsenide, which has a relatively large stable region, but only towards the As-rich compositions^74^. The corresponding elemental maps of Cu and As of the 3D reconstructed volume are displayed in Figure 4d. The distribution of the Cu is not fully homogeneous, forming 5–10 nm clusters distributed throughout the reconstructed volume, which can be explained by the a slight supersaturation of Cu (Cu_3.1-3.4_As) based on the estimated average composition of the particle.

To investigate the 3D morphology of the Cu_3_As particles throughout the enamel, SEM-FIB tomography was performed in the outer enamel and near to the EDJ, Figure 4d. Based on the contrast between the Cu_3_As particles and the HAP matrix, a set of isocontrast surfaces is used to visualize the morphology of the particles (see Methods). Their aspect ratio and oblateness were calculated based on Ref.^75^, and are plotted alongside to quantify the morphology distribution. Close to the EDJ, the particles are larger and more spherical in shape, whereas in the outer enamel region, the particles are finer and more needle-shaped. These observations suggest a strong correlation between the morphology of the Cu_3_As particles and the alignment and size of the HAP grains. The morphology differences of these particles can be ascribed to their growth between the HAP grains, which depends on the HAP grain size and shape.

## Discussion

Our APT results have revealed a high density of organic/carbonate-, and Na-rich clusters at HAP grain boundaries as well as within the core of grains, with a relatively higher density near the EDJ region. Considering these as residual degradation products of amelogenin and non-amelogenins^66^, this could explain the larger grains shown in Figure 1e. This leads us to hypothesise that these clusters contribute or even facilitate the fast growth of the HAP grains^76^ The HAP grains originate from the organic matrix, with a growth mainly occuring on the basal plane (0001). However, it has been shown that incorporation of different cations and anions into HAP grains can influence their growth mechanisms via stabilization of e.g. Mg-rich amorphous calcium phosphate (Mg-ACP) and their morphology. For instance, CO_3_^2-^ ions cause lattice mismatch and hinder active growth directions, which can lead to isotropic growth from all sides of these grains^77,78^. It may not be surprising that some end up trapped within the HAP grains, as discussed in other biomineralization case such as calcite in sea urchin^79^, where however in this case the carbonate (CO_3_^2-^) is the major anion in the lattice. If these are effective at facilitating the nucleation of HAP, their high density could explain the fast growth of the enamel that is necessary for a quick tooth replacement as reported for sauropods ^30^.

A unique feature is the presence of the intergranular Cu_3_As particles in the enamel of this dinosaur tooth. Although metals can be present in enamel in the form of iron-rich coatings^7^, such copper arsenide phases had never been reported in any modern or fossil tooth. These particles are located within the gaps between HAP grains and the morphology is linked to that of the HAP grains.

These could originate from diagenesis, with incorporation of Cu and As from the ambient taphonomic setting into the intergranular phase. Copper and arsenic may originate from the surrounding bedrock or sediment in which the teeth were fossilized. Local reports shows the presence of these elements in the local soil^80^, which could support such an interpretation. Based on the reported diffusivity of Cu nanoparticles in dentine tubules^81^, the diffusion depth can be estimated to be 0.7 mm per month. Therefore, diffusion should have permeated the entire dentine within a few years, especially given that the diffusion of Cu ions is expected to be much faster than that of Cu nanoparticles. It could occur over geological time scales since the age of the dinosaur tooth from the middle or upper dinosaur member of the Tendaguru Formation is stratigraphically constraint to around 150 million years^82^. However, uptake of ionic species from fluid should also result in a higher Cu and As content within the porous dentine, exposed either through the root or through cracks, and the EDS maps (Figure S10) and the APT analysis of the dentine structure show no measurable content of Cu or As (Figure S12).

The shallow burial in a strata only up to 430 m^83^ of the Tendaguru beds and a normal geothermal gradient of 3 °C/100 m both imply that the burial temperature barely exceeded 100 °C and pressures of several hundred bars are not unusual (up to 3 kbar), depending on the depth and exact location of the fossil^84,85^. This was also confirmed by enamel clumped isotope data from the same tooth; the isotopologue thermometry reconstruction of this and other teeth from Tendaguru clearly argue against a deep burial and high temperature exposure of the dinosaur beds^36^. This excludes any high-temperature mechanisms for the formation of these particles^86–89^ or an effect of the temperature and pressure facilitating indiffusion into the enamel.

Another observation speaks against these arisig from diagenesis is that the distribution the Cu_3_As particles within the dental tissues does not show preferential enrichment at the enamel surface, along cracks, or within dentinal tubules, which would be expected if these elements had been introduced primarily through fluid-mediated diagenetic infiltration ^34,90,91^. The absence of such localized enrichment therefore provides additional evidence against extensive secondary infiltration, although it cannot by itself exclude other forms of diagenetic alteration. In addition, Enamel is highly mineralized and densely packed, with very narrow/intermittent intercrystalline spaces – estimated to be in the range of 2 – 6 nm ^39^, so bulk fluid diffusion through intact enamel is strongly restricted^90,92^, unless these fluids were to dissolve part of the HAP enamel crystals.

An alternative hypothesis is the particles formed as the tooth was growing and mineralizing. Studies show that As has a tendency to replace phosphate ion (PO_4_^3-^) in HAP at room temperature, delay the crystallization and expand the grain lattice parameter^93,94^, similar to Mg replacing Ca and stabilizing ACP. Our APT result in Figure 3 shows the Mg ions segregation at grain boundaries. Assuming that As ions were also pushed away toward the edge of the grains during the prematuration process, where there were still some gaps between grains. The Cu could originate from external sources such as plant matter or water consumed during tooth formation^80^. The von Ebner lines suggest that *Giraffatitan brancai* had a rapid tooth replacement rate^30^, which is related to the feeding behavior not involving much oral processing of food to ensure high food intake to meet daily energy requirements to support the giant bodymass of *Giraffatitan brancai*. One could hence assume that the As ejected by the crystallization of the HAP would bond with Cu ion incorporated in the grain boundaries through fluids, forming the observed particles. Figure S13 provides the bright field TEM image and the inverse pole figure of Cu_3_As particles by using transmission kikuchi diffraction in the SEM. The results show that the particles are polycrystalline and in direct contact with the surrounding HAP grains, suggesting that each particle is formed by the merging of multiple growing grains of Cu_3_As, each growing from the surface of a growing or crystallizing HAP grain.

## Conclusion

In this study, we present a multiscale analysis of enamel from the micro- to the nano- and sub-atomic level to understand the structure and composition of a fossil tooth of the sauropod dinosaur *Giraffatitan brancai* using a combined approach of different microscopy techniques, including SEM, SEM-FIB tomography, TEM, and APT. Due to the preservation of the specimen, we were able to identify nanoscale structural features and trace element distributions in the enamel, which provide new insights into both the initial biomineralization mechanism and the postmortem fossilization processes. Key aspects of enamel formation in dinosaurs appear to follow the same biological mechanisms as within extant tetrapods. The widespread appearance of Na- and organic-rich clusters both at the grain boundaries and within HAP grains suggest that they likely acted as nucleation sites which is also supported by positive correlation between the organic-rich cluster density and grain width, contributing to the rapid enamel growth consistent with the short tooth replacement interval inferred from von Ebner line spacing. In addition, we identify copper-arsenide (Cu_3_As) particles distributed throughout the entire enamel structure, a mineral phase so far unknown in any tooth structure, which we hypothesise to have formed during initial tooth mineralization. Our findings highlight the tremendous potential of APT to map the composition of biominerals at near-atomic scales to complementing studies of the biomineralization processes and diagenetic changes of fossils, opening new research avenues for paleontology and the study of hard tissues.

## Acknowledgments

Y.-L. H., T. M. S., A. T., M.D. and B.G are grateful for the financial support from German Research Foundation (DFG) through the award of Leibniz Prize 2020 from B.G. T. M. S. gratefully acknowledges the financial support of the Walter Benjamin Program of the German Research Foundation (DFG) (Project No. 551061178). X.C. gratefully acknowledge the Collaborative Research Centre/Transregio (CRC/TRR) 270 HoMMage-Z01 project funded by the Deutsche Forschungsgemeinschaft (DFG). K. J. is grateful for financial support from the National Research Foundation of Korea (NRF) (grant numbers RS-2025-02483025). B. R. is grateful for the financial support of a Minerva Stiftung Fellowship and Alexander von Humboldt Fellowship. T.T. acknowledges funding by the German National Science Foundation (DFG) Grant TU 148/2-1 for the Emmy Noether-Group “Bone Geochemistry”. This is a contribution to the former DFG Research Unit 533 “Biology of the Sauropod Dinosaurs: The Evolution of Gigantism”.

The authors thank Berlin Naturkundemuseum for providing the dinosaur samples used in this study. The authors are grateful to Uwe Tezins, Katja Angenendt and Andreas Sturm for their support to the APT, SEM, and FIB facilities. Thank Volker Kree for their support to the TEM facilities, Benjamin Breitbach for the XRD and Petra Ebbinghausen for the FTIR measurements at the Max Planck Institute for Sustainable Materials. We would like to thank Stephanie Lohmeier and Ulf Hemmerling from Clausthal University of Technology for providing the FAP samples.

## Data availability

All data needed to evaluate the conclusions in the paper are included in the paper and/or supplementary materials. Additional data for this work can be requested from the contributed authors.

## Supplementary information

Supplementary information is available for this paper.

## Materials and Methods

### Materials and sample preparation

We investigated a well-preserved 150-million-year-old fossil dinosaur tooth of the sauropod *Giraffatitan brancai* from the Late Jurassic Tendaguru Formation^82^. This sauropod fossil was excavated from Tendaguru, Tanzania, during the expedition of German researchers lead by paleontologist Wilhelm von Branca in 1909‒1913^27^. The tooth was cut (Figure S1) and embedded into epoxy resin (Epothin, Buehler, Lake Bluff, IL, USA), followed by grinding (800, 2500, and 4000 grit SiC films; Buehler, Lake Bluff, IL, USA) and polishing with a nap polishing cloth soaked in diamond suspension (3 and 1 µm; Struers Inc., Cleveland, OH, USA) to create a flat surface and remove a possible damaged or oxidized surface layer. After mounted on a stub for microscopical observations, gold coating with a thickness around 15 nm was applied to improve the electric conductivity.

As a compositional reference material for the dinosaur tooth enamel structure, APT specimens of hydroxylapatite (HAP) and fluorapatite (FAP) were prepared and analyzed to identify distinctive fingerprint signals for each system. These fingerprint signals would allow an unambiguous interpretation of the signals in the mass spectra of the dinosaur enamel sample. For the analysis of the reference samples, a sintered HAP sample and a geological FAP grain were used. Further information on sample preparation for the sintered HAP and the origin of the FAP grains can be found below.

#### Hydroxylapatite

For successful APT measurements of the HAP reference, a dense material is necessary due to the particle size and the fact that the synthesized particles are not completely dense it was not possible to prepare an APT specimen directly from powder – instead, a sintered HAP sample was produced. The sintered HAP reference sample was prepared by spark plasma sintering (SPS) (FCT HP D 5), using commercial HAP nano powder with a particle size of <200 nm purchased from Sigma Aldrich (CAS-Nr. 12167-74-7). 3 g of the nano powder was poured in a 20 mm diameter graphite mold lined with graphite foils. Densification was carried out with a heating profile of 50 °C/min up to 800 °C followed by 25 °C/min up to 900 °C where it was held isothermally for 10 min before being cooled down at 25 °C/min with an intermediate holding step at 800 °C for 5 min. The temperature was monitored by a pyrometer via a non-through hole in the top punch. An uniaxial pressure of 38 MPa was applied prior to heating and then monotonously increased up to 63 MPa during the initial heating stage, finally the pressure was relieved during the first cooling stage down to 800 °C. The obtained disc was then extracted, grounded, and polished using 3 µm diamond slurry.

XRD analysis was performed on the initial nano powders and the SPS sample to exclude any possible decomposition during sintering. Apart from the crystalline hexagonal hydroxylapatite signals, only a negligible amount of tricalcium phosphate was observed. The transparency of the SPS sample (Figure S14a) also indicates that the HAP phase did not decompose during the SPS process and confirms the high chemical purity of the sample. The average grain size of the polycrystalline sintered sample is approximately 100–150 nm, calculated from the intergranular fracture surface (Figure S14b). FTIR-ATR measurements of the original nano powder and the SPS sample (Figure S15) confirm the presence of liberational and stretching OH−bands at 629 and 3572 cm^−1^, with no broadening of the 1200–950 cm^−1^ band, which would correspond to tetracalcium phosphate (TTCP) and α-tricalcium phosphate (α-TCP)^95^. Under the parameters used above, SPS enables the densification of hydroxylapatite below its decomposition temperature, without any noticeable phase transitions^96,97^.

#### Fluorapatite

FAP samples are from the mineralogical collection (GeoSammlung) of Clausthal University of Technology. The grains were collected during field trips and are from the Cerro Mercado in the Durango Region in Mexico. Handpicked, 2–3 cm long yellow to slightly brownish apatite grains were embedded in epoxy resin and afterwards polished down to obtain a plane surface. The final polished section was carbon coated. Samples were checked by electron microprobe for fluorine content using energy-dispersive X-ray spectroscopy identifying them as fluorapatite. Electron microprobe analysis of fluorapatite from the Durango region was carried out using a Cameca SX Five field-emission microprobe, housed at Clausthal University of Technology, Institute of Geotechnology and Mineral Resources, Department of Geosciences (Germany). This analytical setting was tested on reference fluorapatite-dispersive mode to minimize diffusion problems and beam-induced damage. The following X-ray lines, spectrometer grains and reference materials (in parentheses) have been used: Ca Kα PET (apatite), Cl Kα LPET (tugtupite), F Kα LPC1 (apatite) and P Kα PET (GaP). Peak measuring time were 30 s for P and 20 s for Ca and Cl, and 10 s for F; one-side background measuring time were half peak measuring time. Background measuring was as follows: P (-/500), Ca (-/550), F (-2600/4000), Cl (-/500). This analytical setting was tested on reference fluorapatite, the measurement of which gave no detectable X-ray line interference that could have influenced the output of quantitative measurement. The result shows 3.0 at.% of F, 23.6 at.% of Ca, 13.9 at.% of P, 0.3 at.% of Cl, and 59.2 at.% of O.

### X-ray diffraction

X-ray diffraction (XRD) was performed using a Rigaku SmartLab 9 kW diffractometer, with a Cu Kα (λ = 1.54059 Å) operated at 45 kV and 200 mA. The scanning range was 5°–120° 2θ with a sampling step of 0.01° at a speed of 1°/min. The obtained patterns were analyzed using MDI JADE 10 software with PDF-4+ database.

### Fourier-transform infrared spectroscopy

FTIR spectra were recorded using an external reflection unit equipped with an ATR sample holder (Seagull, Harrick Scientific Products, Inc., New York) in a Bruker Vertex 70V FTIR spectrometer at an incident angle of 30°. A deuterated triglycine sulphate (DTGS) detector and a KBr beam splitter were used.

For the sintered hydroxylapatite (HAP), the connected Bruker Hyperion 3000 IR microscope with an ATR objective (Ge grain, pressure level approximately 0.5 Newton) and a Mercury Cadmium Telluride (MCT) detector was used.

Each spectrum was recorded with an accumulation of 64 scans and a resolution of 4 cm⁻¹. In ATR mode, the reference spectrum is a single-channel spectrum of the ATR grain.

### Scanning electron microscopy

A scanning electron microscope (Zeiss Sigma 500) equipped with EDAX APEX advanced EDS detector was operated at 15 kV to obtain the overview morphology and element distribution of the specimen. The fracture surface microstructure of the sintered synthetic hydroxylapatite ceramic was examined by high-resolution SEM using a Zeiss Merlin operated at 30 kV and 2 nA. To mitigate charging, the fracture surface was carbon coated (10 nm) using a Gatan PECS II system.

### Transmission electron microscopy sample preparation

A dual-beam scanning electron microscope – focused ion beam (SEM-FIB, Thermo Fisher Instruments, Helios 600i) was used to prepare the TEM specimens at room temperature by using a standard lift-out protocol for TEM lamella preparation^98^. First, Pt was deposited on a selected region-of-interested (ROI) with first electron-beam and followed by the ion-beam deposition using the gas injection system (GIS) to protect the surface of the specimen. Second, the lamella was extracted using the micromanipulator and then mounted on the copper grid for further thinning. Third, the stage was tilted to 52° ± 1.3° on both sides to thin the lamella by decreasing the current step by step until the lamella turned transparent at an electron accelerating voltage of 3 kV. Finally, the stage was tilted to the maximum angle and a rectangle pattern at 5 kV, 15 pA was performed to remove any damaged layers by the ion beam. The thickness of the final TEM lamella was around 60 nm.

### Transmission electron microscopy analysis

The transmission electron microscopy (TEM) images were obtained by using scanning transmission electron microscopy (STEM) mode at an accelerating voltage of 300 kV, camera length of 100 mm, beam convergence angle of 27 mrad and a high-angle annular dark field (HAADF) detector was used with a collection angle range of 78–200 mrad on a probe-corrected Thermo Fisher Titan Themis G2 microscope (pc-TEM). Images and selected area electron diffraction (SAED) patterns were obtained on a CMOS (metal-oxide-semiconductor) camera (4k×4k pixels). TEM bright field images were obtained using Jeol-JEM2200FS at an accelerating voltage of 80 kV with an omega energy filter inserted.

We employed 4D-STEM to map crystallographic orientations and phase distributions using a JEM-2200FS microscope (JEOL Ltd.) operated at 200 kV and equipped with a TVIPS camera for recording scanning nanobeam diffraction patterns. The same needle-shaped specimen was subsequently analyzed by APT, enabling direct, site-specific correlation of crystal structure and chemical composition at the atomic scale.

### Atom probe tomography specimen preparation

APT specimen preparation was performed, and a series of needle-shaped specimens were prepared in the other SEM-FIB (Thermo Fisher Instruments, Helio 5CX) at room temperature, using a standard lift-out protocol^99^ from the selected ROI. First, Pt was deposited in the electron-beam and ion-beam using the GIS to protect the surface of ROI and prevent any structural changes to the internal material. Three trenches were cut into the material before the lamella was lifted out using the micromanipulator and mounted with Pt to the silicon support post (microtip array from Cameca Inc.) using the GIS. Following, by annular milling, whereby the ion current and inner diameter of the annular pattern are decreased until the tip radius is less than 100 nm (Figure S2). Finally, a low accelerating voltage and low current at 5 kV and 7 pA were applied to remove the beam damage on the surface of the tip. APT specimens were prepared in different areas of the enamel structure, but always at the same horizonal line perpendicular to the enamel dentine junction (EDJ) toward the edge of the enamel structure, to minimize any structural changes throughout different areas.

### Atom probe tomography measurement, data reconstruction and processing

APT measurements were carried out using Cameca LEAP 5000 XR, with a base temperature of 40 K, a target detection rate of 0.5 % and a laser repetition frequency of 125 kHz. Laser energy ranging from 100 pJ to 150 pJ for tooth enamel (six datasets collected over 10 mio. events in total 12 runs), sintered HAP (two datasets collected over 10 mio. events in total three runs), and FAP samples (two datasets collected over 10 mio. events in total three runs). For tooth dentine (two datasets collected over 20 mio. events in total three runs) and Cu_3_As particles (two datasets collected over two mio. events), a laser energy between 10 and 25 pJ was used.

Data reconstruction and post-processing were done using the software AP Suite 6.3 by Cameca Inc.. For all reconstructions, the specimen radius evolution model was used, based on the assumption of a constant shank angle^100^. The initial radius and shaft angle were determined from the SEM images. A field factor of k*_f_* = 3.3 and an image compression factor of ξ = 1.65 were applied to all reconstructions. All identified signals for the tooth enamel structure, HAP, and FAP specimens are listed in Table S1.

Mass spectra of HAP, FAP, and the enamel are plotted in Figure S12. All peaks have been normalized to the highest peak. HAP, FAP and tooth enamel are mainly comprised of the combination of Ca, P, O, and H. Signals of Mg^2+^, CaF^+^ and P*_x_*O*_y_*F*_z_*^1–3+^ are only present in the mass spectra of FAP and tooth enamel, where the Mg^2+^ in the FAP originates from the geological origin of the sample. In addition, the signal at 19 Da, which can be assigned to either H_3_O^+^ or F^+^, is only prominent in the HAP and tooth enamel mass spectra, but not in the FAP mass spectra. This suggests that the signal at 19 Da can be unambiguously assigned to H_3_O^+^. It also indicates that it is almost impossible to ionize and evaporate an F ion due to its high ionization energy (17.423 eV)^101^. Therefore, only signals containing F ion could be observed in the form of compounds bonded with Ca^2+^ or phosphate groups.

### SEM-FIB tomography

SEM-FIB tomography was performed on Helios 5CX (Ga) SEM-FIB instrument from Thermo Fisher Scientifc. First, a thin layer of ion beam Pt was deposited on the surface of the ROI using the GIS inside the SEM-FIB instrument, with an acceleration voltage of 30 kV and a current of 0.23/0.43 nA. Second, a Pt marker was deposited on the surface near the ROI for the Auto Slice and view mode. Afterwards, three trenches were cut around the ROI (left, right, and front) to avoid shadowing effects during auto slice and view mode. Finally, the FIB Tomography acquisition was acquired in Auto Slice and View mode, with an ion accelerating voltage of 30 kV, an ion current of 0.23 nA, and a slice thickness of 20 nm per pass. Electron beam images were required using an electron accelerating voltage of 5 kV, a probe current of 86 pA, and a dwell time of 3 µs. The acquired images were first segmented in ilastik 1.4.1 using pixel classification method^102^. After segmented, all the images were imported into Dragonfly 3D World 2024.1 to perform the 3D volumetric reconstruct ions.

