## Supplementary material for "Nanoscale chemical tomography reveals organic and inorganic clusters in fossilised dinosaur tooth"

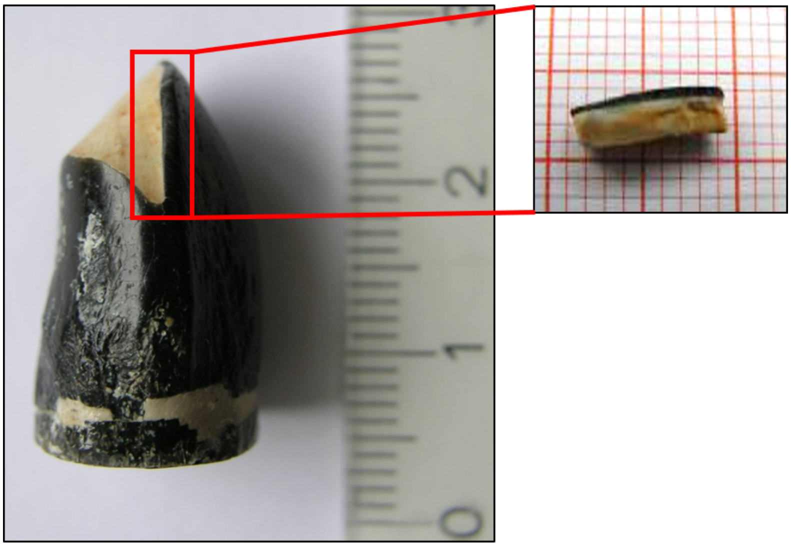

**Figure S1** Fossil tooth sample (MB.R.2181) of the Late Jurassic sauropod *Giraffatitan brancai* from Tendaguru. The black outer sub-mm layer is enamel and the yellowish material in the center is dentine. A piece containing both dental tissues was cut for chemical and structural analysis from the top part of the tooth. (Modified from[1])

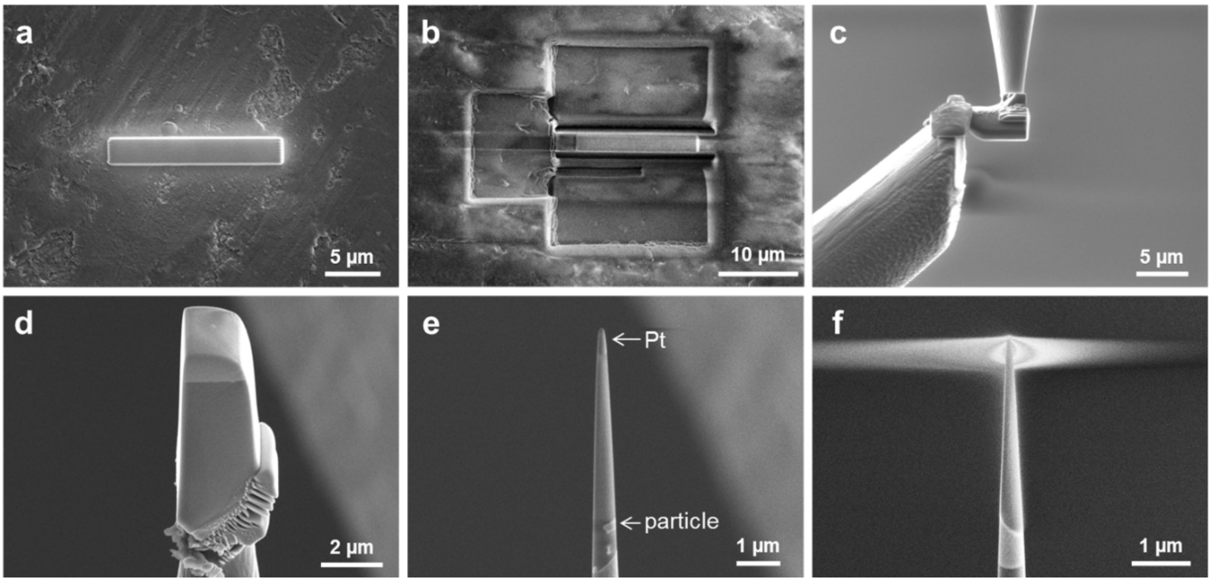


**Figure S2** APT specimen preparation using the 52° lift-out method. (a) The dinosaur enamel structure with Pt deposition on the selected area. (b) Three trenches were cut into the specimen for further lamella extraction. (c) The lamella was glued onto a commercial Si post. (d) The mounted lamella before the milling process. (e) Annular milling was performed, and (f) the final specimens shaped as sharp needles with a tip radius under 100 nm.

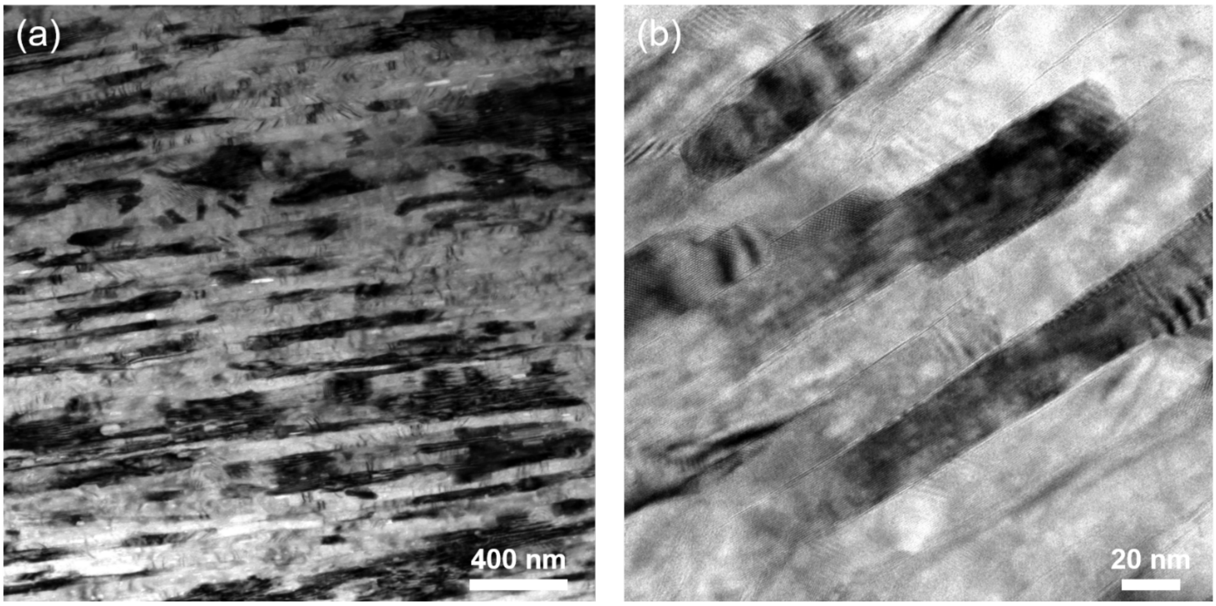


**Figure S3** TEM bright field images: (a) *Giraffatitan brancai* enamel (around the middle region) HAP grains. (b) Higher magnification image of some HAP grains observed in another area of (a).

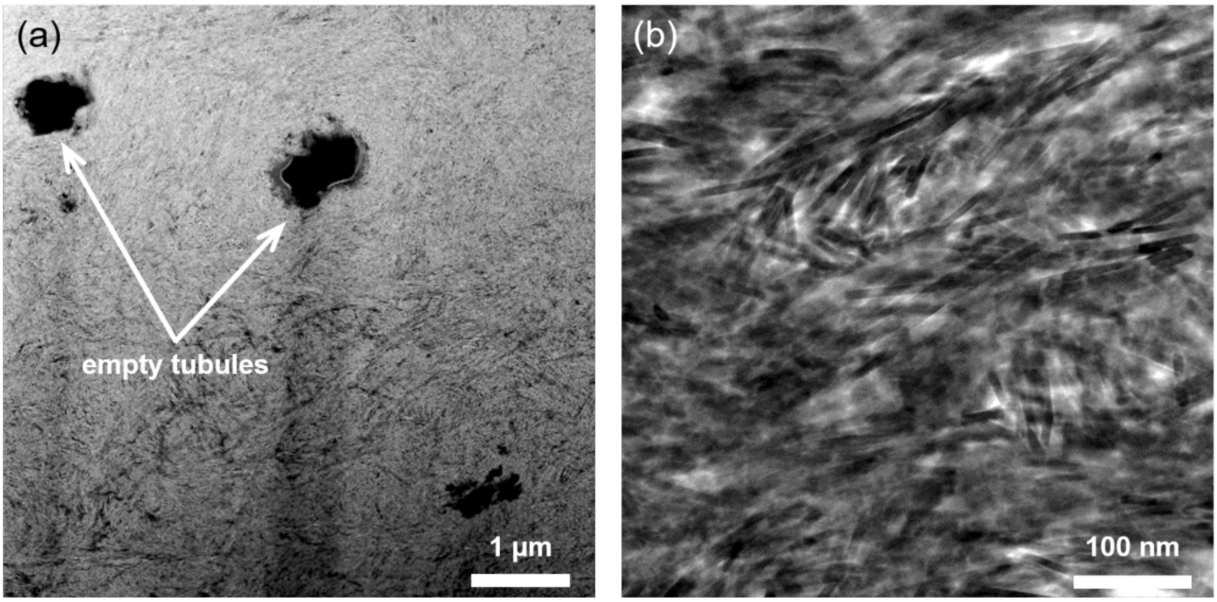


**Figure S4** TEM bright field images: (a) Dentine of the *Giraffatitan brancai* tooth showing cross sections of the empty dentinal tubules suggesting a good preservation (i.e., no secondary mineral or sediment infills). (b) Higher magnification image of needle-shaped HAP dentine grains.

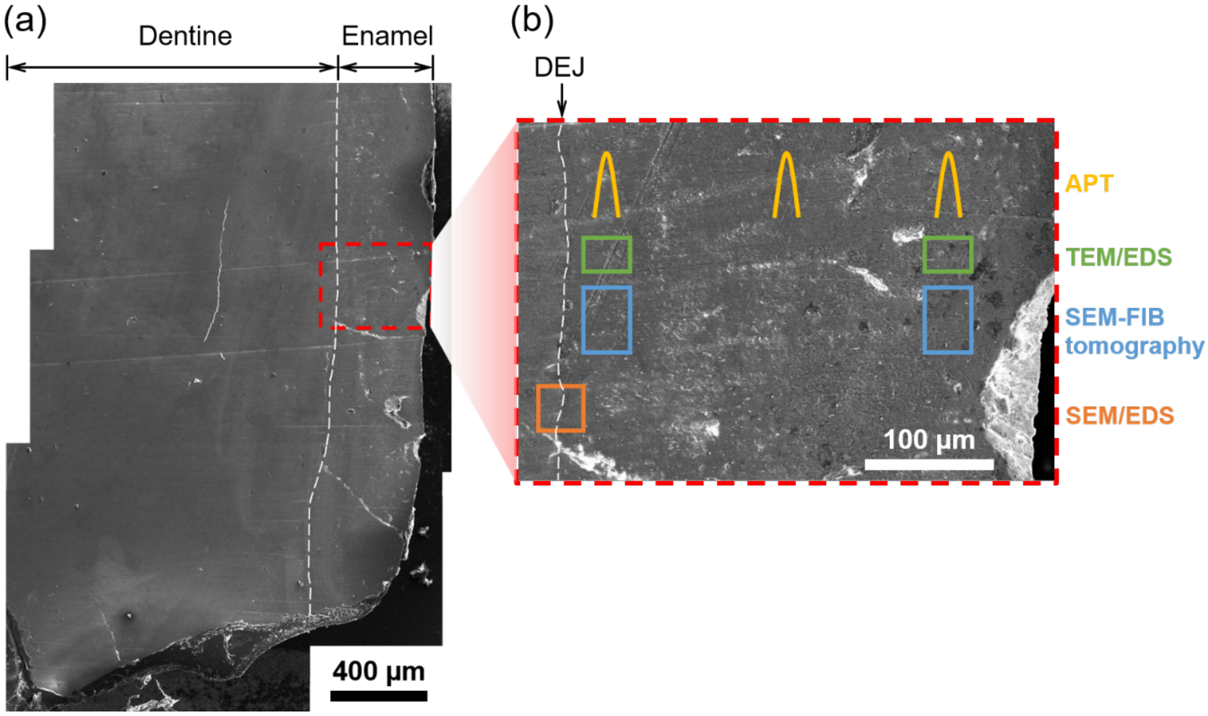


**Figure S5** SE image of the polished sauropod tooth section with the dentin enamel junction (DEJ). In the enlarged image of the enamel cross section the locations are schematically shown where nm-sized subsamples were prepared for subsequent chemical and structural analysis using FIB-SEM, TEM, SEM and APT.

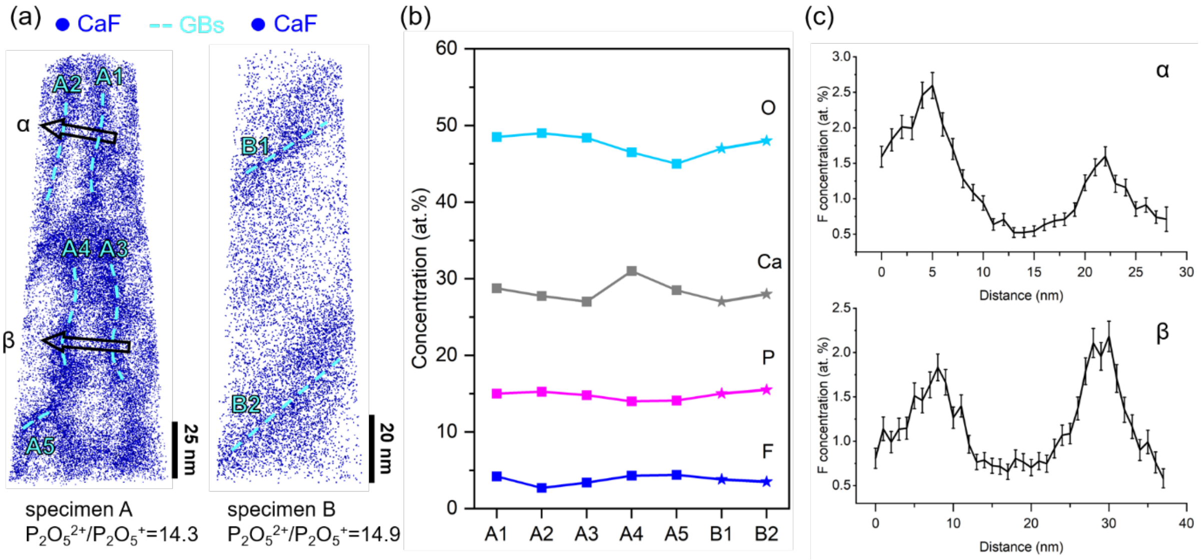


**Figure S6** (a) APT 3D reconstructions of two *Giraffatitan brancai* enamel conical subsamples lift-out in the central part of the enamel (see **Figure S5**). Sky-blue dashed lines mark the grain boundaries, and the arrows indicate the directions of 1D concentration profile. (b) Comparison of major element compositions at grain boundaries. (c) 1D F-concentration profiles.

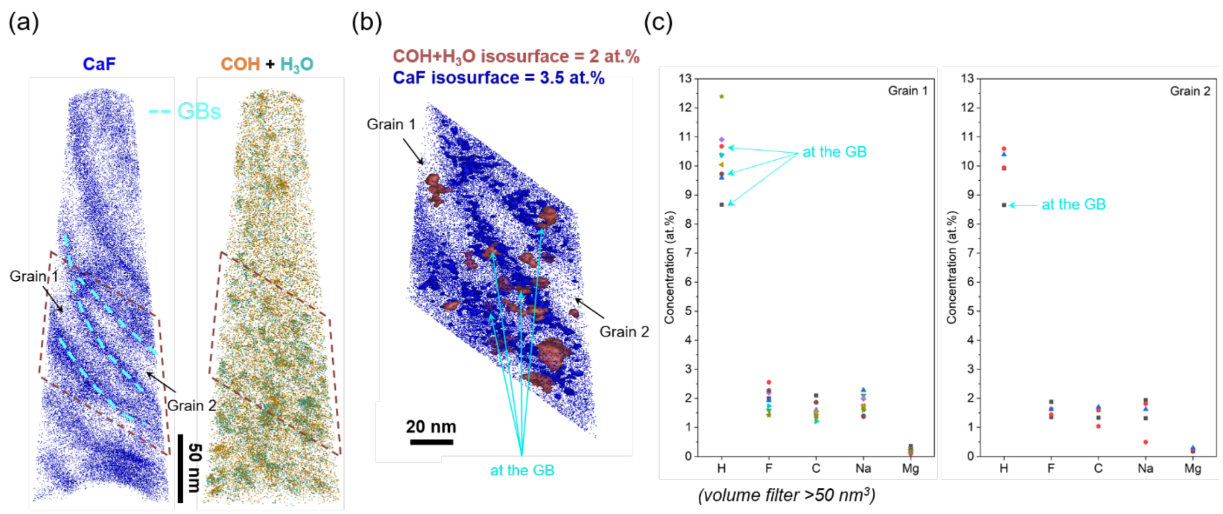


**Figure S7** Cluster composition comparison of the specimen lifted-out near the DEJ (see **Figure S5**).

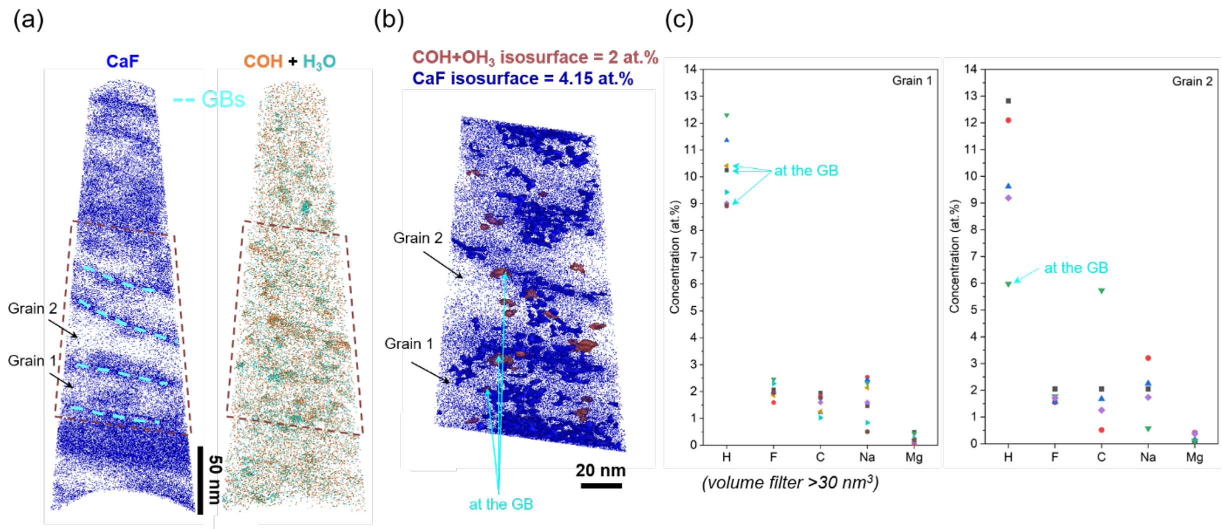


**Figure S8** Cluster composition comparison of the specimen lift-out near the edge of the enamel (see **Figure S5**). In grain 2, the H gap of the cluster at the GB is because this specific cluster consists of more COH and less $H_3O$.

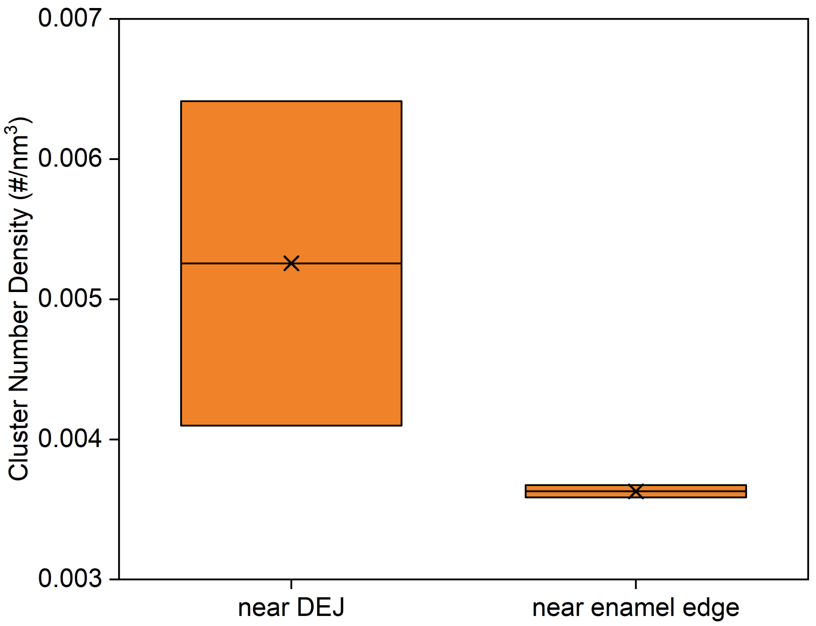


**Figure S9** The cluster number density calculation of COH from regions near the DEJ and near the enamel edge. Two specimens from each region were calculated using cluster analysis function in AP Suite software with the parameter of $d_{max}$=2 nm, order=1 ion, $N_{min}$=10 ions, and L=E=0.1 nm.

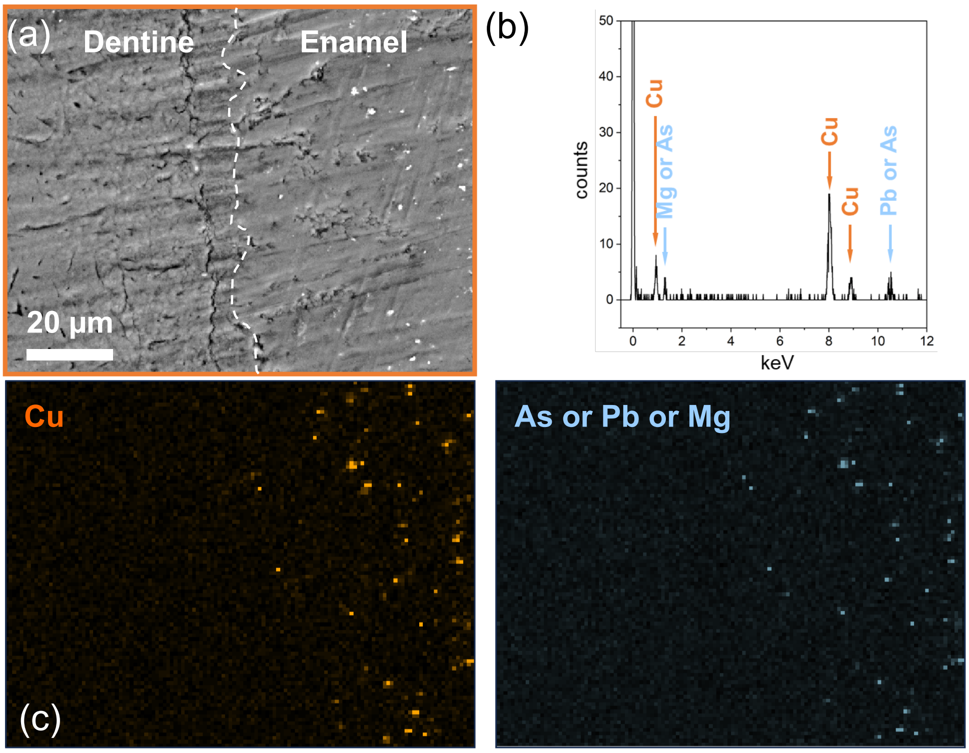


**Figure S10** (a) secondary electron micrograph from Figure 4a, (b) EDS spectrum showing the main peak overlaps and (c) EDS mapping showing colocalization of Cu, and either As, Pb or Mg.

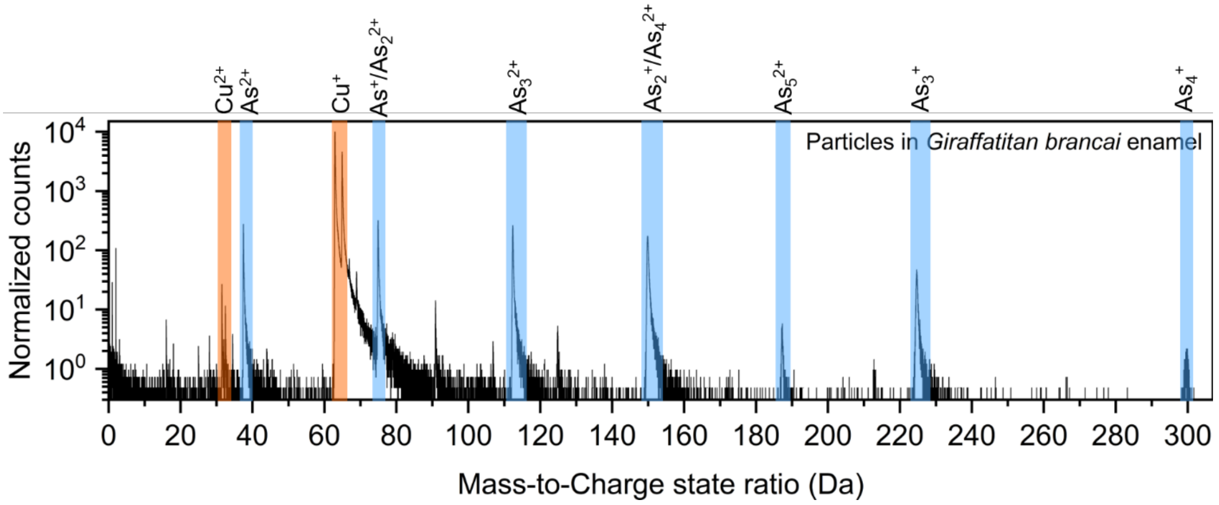


**Figure S11** APT Mass spectrum of the nm-sized particles in *Giraffatitan brancai* enamel structure which indicates that they are composed of copper and arsenic.

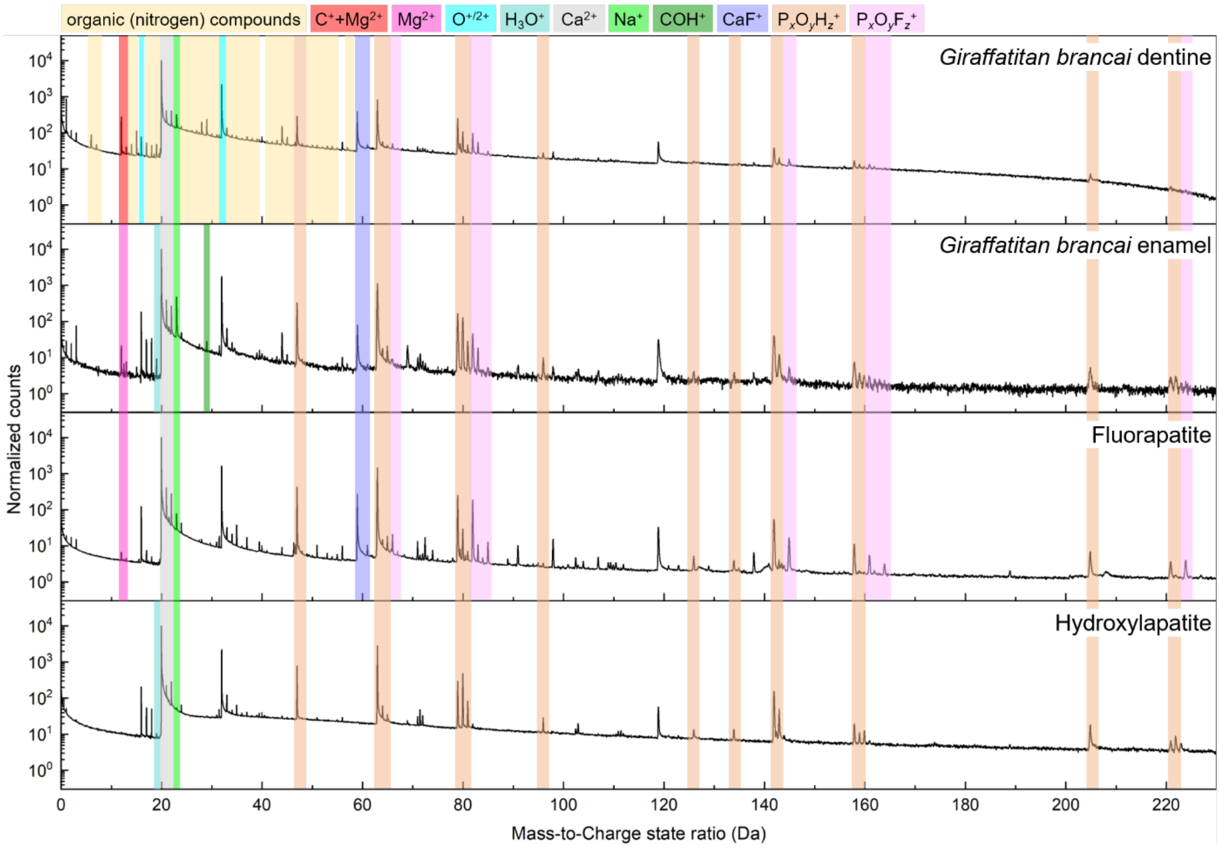


**Figure S12** Mass spectra comparison between *Giraffatitan brancai* dentine, *Giraffatitan brancai* enamel, a magmatic fluorapatite (FAP), and sintered synthethic hydroxylapatite (HAP). All spectra are normalized to the highest peak.

**Table S1** Comparison peak identification between *Giraffatitan brancai* enamel, pure hydroxylapatite (HAP) and fluorapatite (FAP).

| **Mass-to-charge state ratio (Da)** | | **Enamel** | | **Hydroxylapatite** | | **Fluorapatite** | |
|---|---|---|---|---|---|---|---|
| **Lower limit** | **Higher limit** | **Ion/molecule type** | **Charge state** | **Ion/molecule type** | **Charge state** | **Ion/molecule type** | **Charge state** |
| 0.99 | 1.03 | H | + | H | + | H | + |
| 2.00 | 2.03 | $H_2$ | + | $H_2$ | + | $H_2$ | + |
| 3.01 | 3.05 | $H_3$ | + | $H_3$ | + | $H_3$ | + |
| 6.99 | 7.04 | N | 2+ | | | | |
| 11.95 | 12.05 | Mg | 2+ | | | Mg | 2+ |
| 12.46 | 12.56 | Mg | 2+ | | | Mg | 2+ |
| 12.93 | 13.09 | Mg | 2+ | | | Mg | 2+ |
| 14.94 | 15.12 | NH | + | | | | |
| 15.46 | 15.54 | | | | | P | 2+ |
| 15.96 | 16.07 | O | + | O | + | O | + |
| 16.94 | 17.11 | OH | + | OH | + | OH | + |
| 17.95 | 18.09 | $OH_2$ | + | $OH_2$ | + | $OH_2$ | + |
| 18.91 | 19.13 | $OH_3$ | + | $OH_3$ | + | | |
| 19.88 | 20.08 | Ca | 2+ | Ca | 2+ | Ca | 2+ |
| 20.94 | 21.05 | $PO_2$ | 3+ | $PO_2$ | 3+ | $PO_2$ | 3+ |
| 21.44 | 21.52 | Ca | 2+ | Ca | 2+ | Ca | 2+ |
| 21.95 | 22.05 | Ca | 2+ | Ca | 2+ | Ca | 2+ |
| 22.95 | 23.05 | Na | + | Na | + | Na | + |
| 23.93 | 24.01 | Mg | + | Ca | 2+ | Mg | + |
| 27.45 | 27.54 | Mn | + | | | Mn | + |
| 27.85 | 28.11 | CO | + | | | CO | + |
| 28.94 | 29.08 | COH | + | | | | |
| 29.95 | 30.07 | $CO_3$ | 2+ | | | | |
| 30.93 | 31.02 | P | + | P | + | P | + |
| 31.43 | 31.52 | $PO_2$ | 2+ | $PO_2$ | 2+ | $PO_2$ | 2+ |
| 31.92 | 32.09 | $O_2$ | + | $O_2$ | + | $O_2$ | + |
| 32.95 | 33.05 | POF | 2+ | $O_2H$ | + | POF | 2+ |
| 33.95 | 34.06 | $O_2H_2$ | + | $O_2H_2$ | + | $O_2H_2$ | + |
| 34.93 | 35.04 | | | Cl | + | Cl | + |
| 35.90 | 36.06 | $CaO_2$ | 2+ | $CaO_2$ | 2+ | $CaO_2$ | 2+ |
| 36.88 | 37.06 | | | Cl | + | Cl | + |
| 39.43 | 39.52 | $PO_3$ | 2+ | $PO_3$ | 2+ | $PO_3$ | 2+ |
| 39.92 | 40.04 | Ca | + | Ca | + | Ca | + |
| 40.44 | 40.55 | $PO_3H_2$ | 2+ | $PO_3H_2$ | 2+ | | |
| 40.88 | 41.13 | $PO_2F$ | 2+ | | | $PO_2F$ | 2+ |
| 43.92 | 44.05 | $CO_2$ | + | $CO_2$ | + | $CO_2$ | + |
| 44.92 | 45.10 | $CO_2H$ | + | | | | |
| 46.88 | 47.09 | PO | + | PO | + | PO | + |
| 50.90 | 51.03 | | | $O_2F$ | + | $O_2F$ | + |
| 54.89 | 55.03 | $P_2O_3$ | 2+ | | | | |
| 55.91 | 56.05 | CaO | + | CaO | + | CaO | + |
| 58.83 | 59.32 | CaF | + | | | CaF | + |

| | | | | | | | |
|---|---|---|---|---|---|---|---|
| 60.82 | 61.15 | CaF | + | | | CaF | + |
| 62.82 | 63.21 | $PO_2$ | + | $PO_2$ | + | $PO_2$ | + |
| 63.88 | 64.16 | $PO_2H$ | + | $PO_2H$ | + | $PO_2H$ | + |
| 64.89 | 65.10 | $PO_2H_2$ | + | $PO_2H_2$ | + | $PO_2H_2$ | + |
| 65.84 | 66.11 | POF | + | | | POF | + |
| 66.83 | 67.16 | | | | | POFH | + |
| 71.38 | 71.58 | $P_2O_5H$ | 2+ | $P_2O_5H$ | 2+ | $P_2O_5H$ | 2+ |
| 71.87 | 72.07 | $CaO_2$ | + | $CaO_2$ | + | $CaO_2$ | + |
| 72.34 | 72.65 | $P_2O_4F$ | 2+ | | | $P_2O_4F$ | 2+ |
| 73.81 | 74.13 | | | | | $P_2O_3F_2$ | 2+ |
| 78.84 | 79.16 | $PO_3$ | + | $PO_3$ | + | $PO_3$ | + |
| 79.85 | 80.19 | $PO_3H$ | + | $PO_3H$ | + | $PO_3H$ | + |
| 80.83 | 81.16 | $PO_3H_2$ | + | $PO_3H_2$ | + | $PO_3H_2$ | + |
| 81.90 | 82.06 | $PO_2F$ | + | $PO_3H_3$ | + | $PO_2F$ | + |
| 82.81 | 83.17 | $CaP_2O_4$ | 2+ | | | $CaP_2O_4$ | 2+ |
| 83.88 | 84.13 | | | | | $PO_2F$ | + |
| 84.79 | 85.19 | $POF_2$ | + | | | $POF_2$ | + |
| 88.82 | 89.08 | $P_4OF_2$ | 2+ | | | $P_4OF_2$ | 2+ |
| 90.75 | 91.25 | $C_6OH_3$ | + | | | $C_6OH_3$ | + |
| 95.81 | 96.11 | $PO_4H$ | + | $PO_4H$ | + | $PO_4H$ | + |
| 97.73 | 98.22 | $PO_3F$ | + | | | $PO_3F$ | + |
| 100.79 | 101.12 | | | | | $PO_2F_2$ | + |
| 102.31 | 102.59 | $P_3O_7$ | 2+ | $P_3O_7$ | 2+ | $P_3O_7$ | 2+ |
| 102.76 | 103.12 | $P_3O_7H$ | + | $P_3O_7H$ | + | $P_3O_7H$ | + |
| 103.77 | 104.16 | $POF_3$ | + | | | $POF_3$ | + |
| 106.81 | 107.19 | $CaPO_2$ | + | | | $CaPO_2$ | + |
| 110.36 | 110.56 | | | $P_3O_8$ | 2+ | $P_3O_8$ | 2+ |
| 110.84 | 111.05 | | | $P_3O_8H$ | 2+ | | 2+ |
| 111.33 | 111.59 | | | $P_3O_8H_2$ | 2+ | | 2+ |
| 118.72 | 119.35 | $CaPO_3$ | + | $CaPO_3$ | + | $CaPO_3$ | + |
| 122.71 | 123.12 | $POF_4$ | + | | | $POF_4$ | + |
| 125.71 | 126.13 | $P_2O_4$ | + | $P_2O_4$ | + | $P_2O_4$ | + |
| 128.71 | 129.14 | | | | | $P_2O_3F$ | + |
| 133.78 | 134.08 | $P_4O_9$ | 2+ | $P_4O_9$ | 2+ | $P_4O_9$ | 2+ |
| 137.63 | 138.29 | $CaPO_3F$ | + | | | $CaPO_3F$ | + |
| 141.66 | 142.25 | $P_2O_5$ | + | $P_2O_5$ | + | $P_2O_5$ | + |
| 142.69 | 143.25 | $P_2O_5H$ | + | $P_2O_5H$ | + | $P_2O_5H$ | + |
| 144.54 | 145.44 | $P_2O_4F$ | + | | | $P_2O_4F$ | + |
| 157.69 | 158.25 | $P_2O_6$ | + | $P_2O_6$ | + | $P_2O_6$ | + |
| 158.70 | 159.18 | $P_2O_6H$ | + | $P_2O_6H$ | + | | + |
| 159.75 | 160.14 | $P_2O_6H_2$ | + | $P_2O_6H_2$ | + | | + |
| 160.65 | 161.17 | $P_2O_5F$ | + | | | $P_2O_5F$ | + |
| 161.71 | 162.25 | $Ca_2O_5H_2$ | + | | | | |
| 162.67 | 163.24 | $Ca_2O_5H_3$ | + | | | | |
| 163.62 | 164.23 | $P_2O_4F_2$ | + | | | $P_2O_4F_2$ | + |
| 204.51 | 205.28 | $P_3O_7$ | + | $P_3O_7$ | + | $P_3O_7$ | + |
| 220.57 | 221.20 | $P_3O_8$ | + | $P_3O_8$ | + | $P_3O_8$ | + |
| 221.53 | 222.20 | $P_3O_8H$ | + | $P_3O_8H$ | + | | + |
| 222.54 | 223.35 | $P_3O_8H_2$ | + | $P_3O_8H_2$ | + | | + |

| | | | | | | | |
|---|---|---|---|---|---|---|---|
| 223.43 | 224.44 | $P_3O_7F$ | + | | | $P_3O_7F$ | + |
| 283.34 | 284.29 | $P_4O_{10}$ | + | $P_4O_{10}$ | + | $P_4O_{10}$ | + |
| 284.29 | 285.52 | $P_4O_{10}H$ | + | $P_4O_{10}H$ | + | $P_4O_{10}H$ | + |

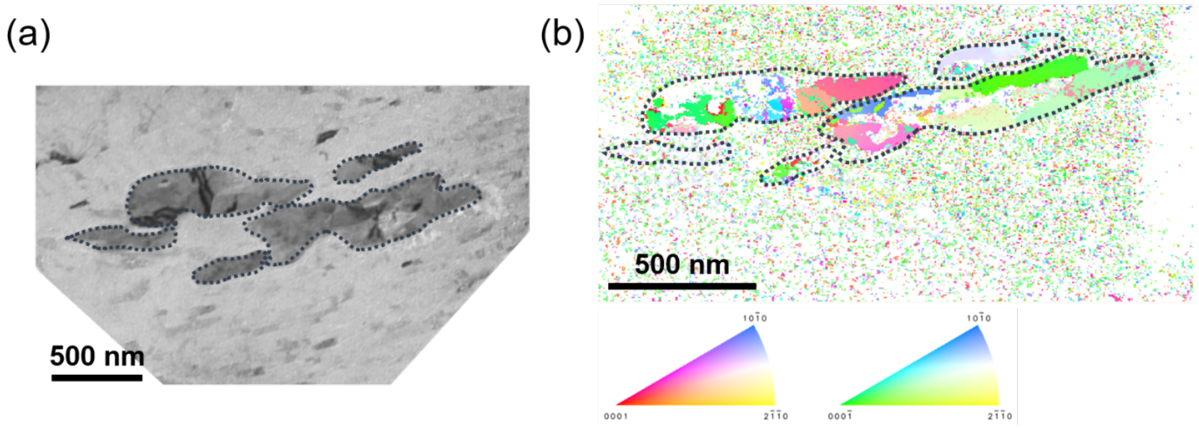


**Figure S13** (a) TEM bright field image of *Giraffatitan brancai* enamel (the orange dashed lines were used to emphasize the particles). (b) Spherical-indexing inverse pole figure image of the particles by using Transmission Kikuchi diffraction (TKD or t-EBSD) technique with acceleration voltage of 20kV, current of 1nA, working distance of 2.6mm, and scanning step size of 5nm. Noted that the image was compressed due to the beam shift during the measurement process.

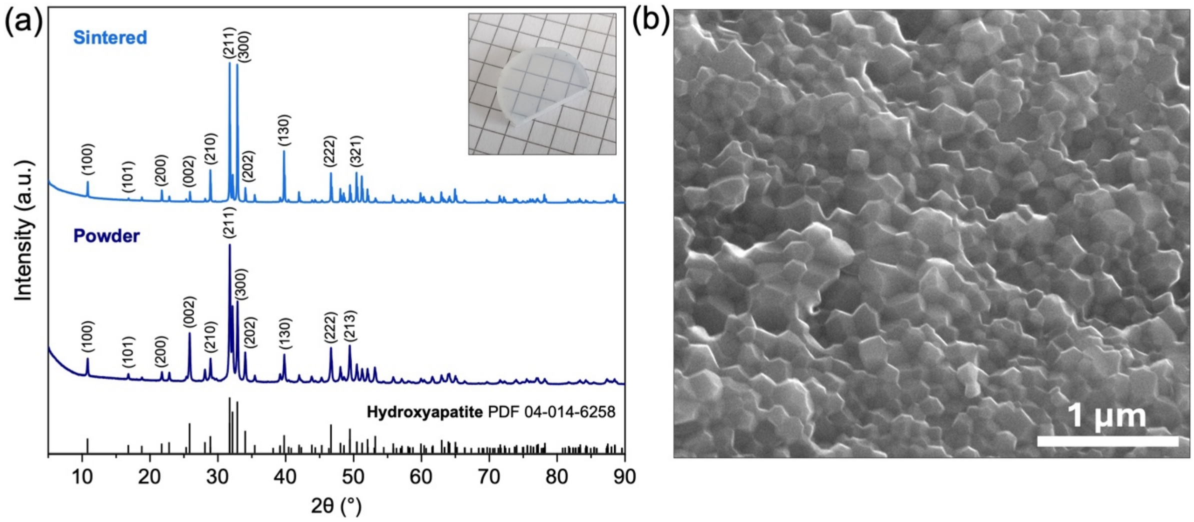


**Figure S14** Sintered hydroxylapatite characterization. (a) XRD plot of initial nanopowder and sintered HAP sample, depicting crystalline hexagonal hydroxylapatite with a negligible amount of tricalcium phosphate (TCP)[2]. The inset shows a photograph of the transparent sintered sample placed on a regular math paper where each square is 5x5 mm$^2$. (b) Intergranular fracture surface of the sintered hydroxylapatite, showcasing the fine grain morphology with the grain average size being approximately 100–150 nm.

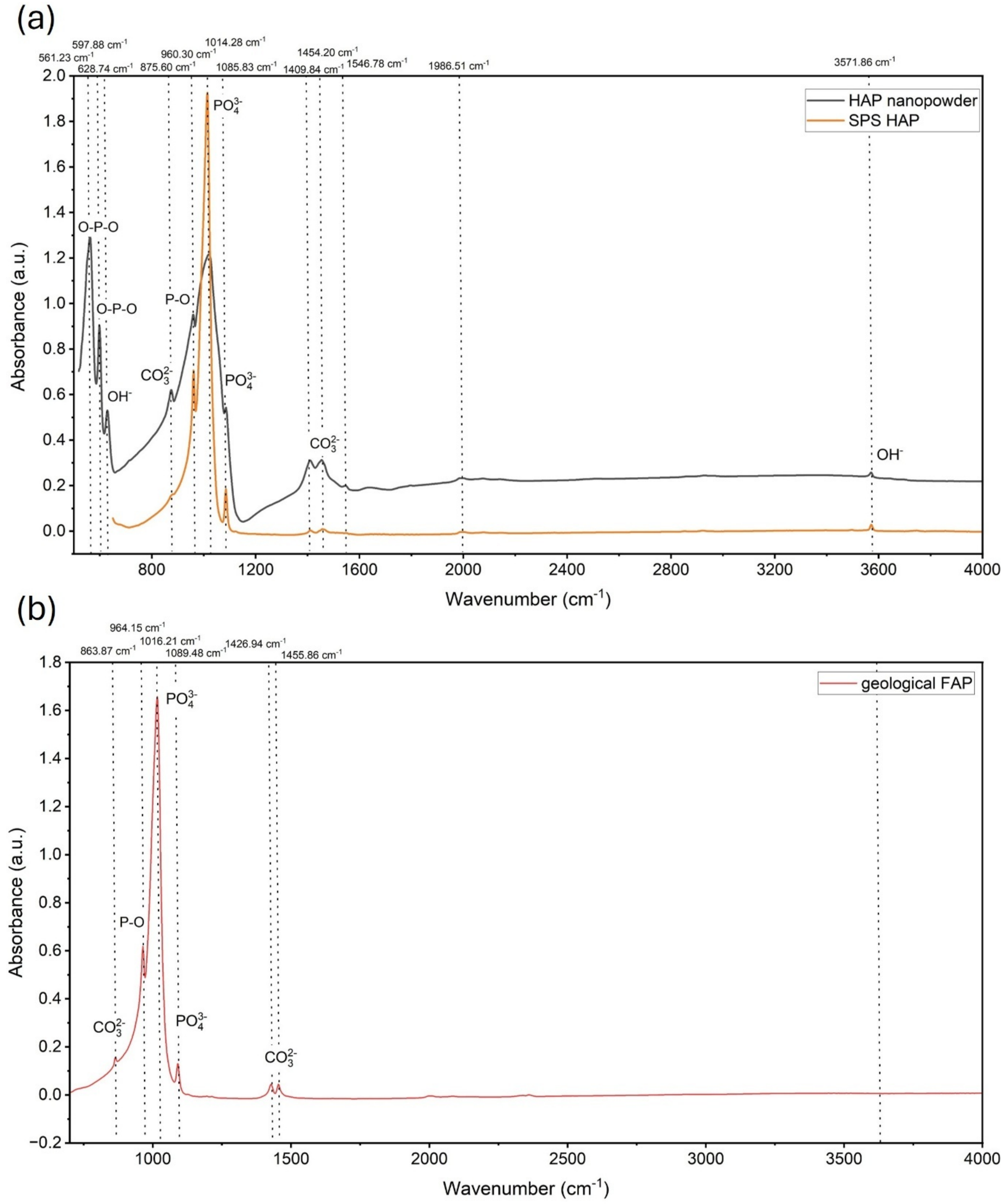


**Figure S15** FTIR spectra of (a) the initial HAP nanopowder and the spark plasma sintered (SPS) HAP sample and (b) the geological FAP sample. $CO_3^{2-}$ peaks reduce and the full width at half maximum (FWHM) of the peak broadens due to the sintering process.
